# ZIP-5/bZIP transcription factor regulation of folate metabolism is critical for aging axon regeneration

**DOI:** 10.1101/727719

**Authors:** Vanisha Lakhina, Melanie McReynolds, Daniel T. Grimes, Joshua D. Rabinowitz, Rebecca D. Burdine, Coleen T. Murphy

## Abstract

Aging is associated with reduced capacity for tissue repair, perhaps the most critical of which is a decline in the ability of aged neurons to recover after injury. Identifying factors that improve the regenerative ability of aging neurons is a prerequisite for therapy design and remains an enormous challenge, yet many of the genes that play a role in regeneration of youthful axons do not regulate axon regeneration in older animals^2,9^, highlighting the need to identify aging-specific regeneration mechanisms. Previously, we found that increased DAF-16/FOXO activity enhances the regenerative ability of mechanosensory axons in aged animals^9^. Here we show that DAF-16/FOXO mediates its pro-regenerative effects by upregulating folate metabolism genes via the ZIP-5 bZIP transcription factor. Remarkably, dietary folic acid supplementation improves the regeneration of aging *C. elegans* axons. Enzymes regulating folate metabolism are also up-regulated in regenerating zebrafish fins, and we show that dietary folic acid supplementation post-amputation enhances fin regrowth in aging zebrafish. Our results demonstrate that boosting folate metabolism is a conserved and non-invasive approach to increase the regenerative capacity of aging neurons and tissues. Given that lower folate status has been linked with reduced cognition in the elderly^17^, maintaining optimal folate metabolism may be a general strategy to achieve healthy brain aging.

---

The FOXO transcription factor plays a conserved role in regulating the rate of neuronal aging. We previously showed that in *C. elegans*, increased DAF-16/FOXO activity in *daf-2* insulin receptor mutants improves memory formation and maintains the ability of PLM mechanosensory neurons to regenerate injured axons well into middle age^9, 11^. Consistent with a role in regulating neuronal aging, young adult FOXO1/3/4 knockout mice display increased axonal degeneration, higher microglial activation, and reduced startle reflex compared to wild-type controls^8^.

To uncover how DAF-16/FOXO improves axon regeneration in aged animals, we profiled isolated neurons to identify neuron-specific FOXO targets^9^ (Fig. 1a; Supplemental Table 1). Neuronal FOXO targets are largely distinct from genes required to regrow injured larval axons in wild-type animals^3,9^ (Fig. 1a), suggesting that DAF-16/FOXO regulates adult axonal regeneration through mechanisms that are distinct from larval axon regeneration. The Dual-Leucine zipper Kinase DLK-1 is the only known DAF-16/FOXO target that regulates both larval and adult regeneration^2,3^, while other DAF-16/FOXO neuronal targets mediate distinct functions such as cellular stress response and amino acid metabolism that are not regulated by larval regeneration genes (Fig. 1b). However, two genes, the C-mannosyltransferase *dpy-19* and the bZIP transcription factor *zip-5*, are FOXO targets (Fig. 1a, Extended Data Fig. 1a, Supplemental Table 1) that function in larval axon regeneration in wild-type animals. We found that ZIP-5 is required for re-growth of injured PLM axons in aging *daf-2* mutants: reducing *zip-5* in either young (Day 1 adults) or middle-aged (Day 5) *daf-2* mutants significantly reduces the repair of PLM axons upon laser microsurgery (Fig. 1c-f), identifying a new role for ZIP-5 in adult axonal regeneration. *zip-5* is not required for DAF-16/FOXO-mediated extension of memory^9^, or for the suppression of age-related morphological defects in neurons (Extended Data Fig. 1b,c), suggesting that DAF-16/FOXO acts through ZIP-5 to selectively enhance axonal regeneration in aging *daf-2* mutants.

**Figure 1.**
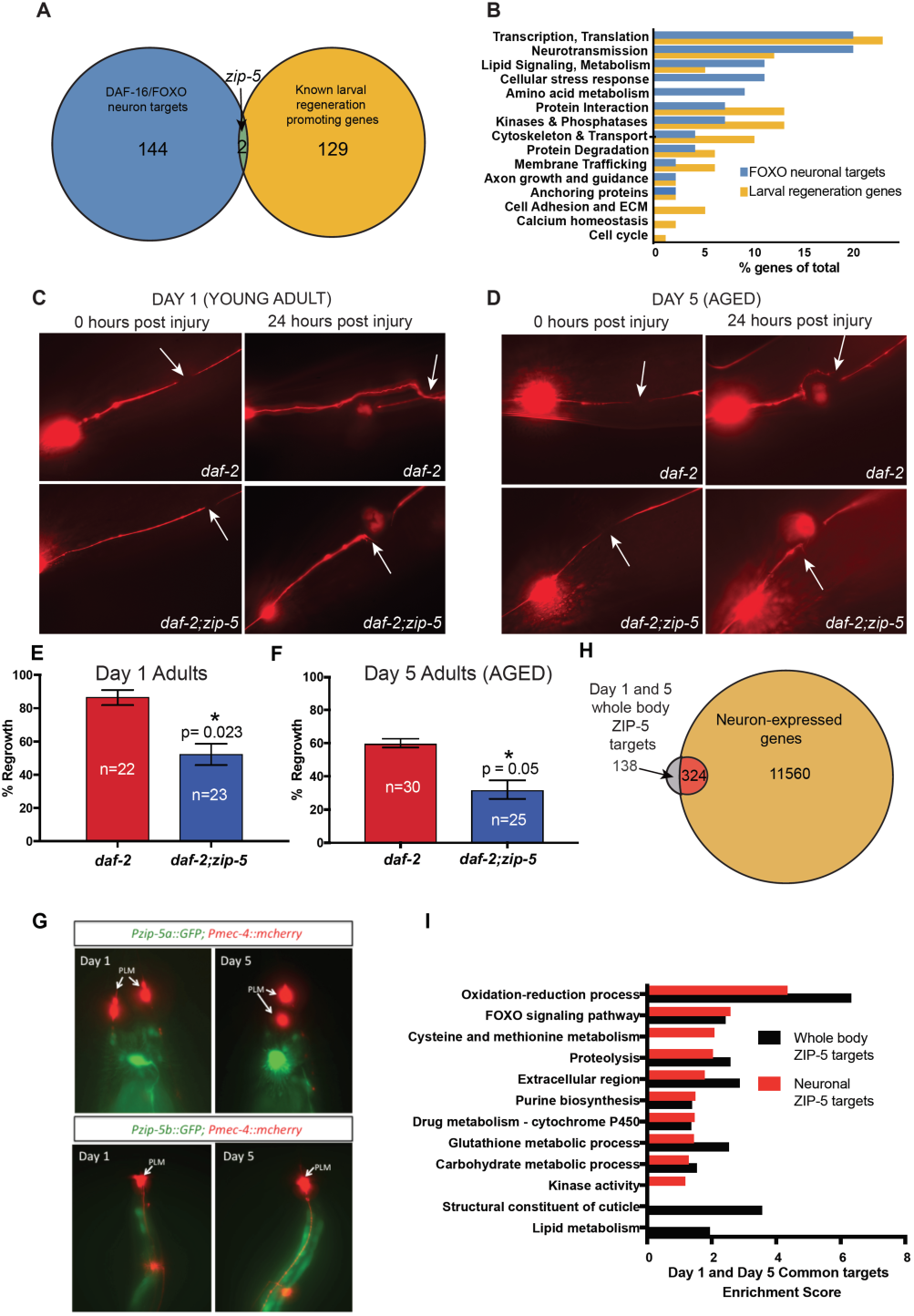
The DAF-16/FOXO target ZIP-5 mediates the improved axonal regeneration of aged *daf-2* mutants by altering metabolism. **a**, *zip-5* is a DAF-16/FOXO target that is required for the regeneration of larval axons. **b**, DAF-16/FOXO neuronal targets and larval regeneration genes mediate overlapping as well as distinct functions. **c, e**, Young day 1 adult *daf-2* mutants regenerate axons at a high frequency, while *daf-2;zip-5* mutants exhibit reduced regeneration. **d, f**, Aged day 5 adult *daf-2;zip-5* mutants have reduced axonal regeneration compared to *daf-2* mutants. **g, *zip-5a*** is expressed in non-PLM neurons, while *zip-5b* is expressed in the intestine in Day 1 and 5 adults. **h**, Microarray analysis comparing *daf-2 vs. daf-2;zip-5* identified 462 genes that are upregulated by ZIP-5 in Day 1 and 5 animals. 324 of the 462 ZIP-5 target genes are expressed in neurons (p-value = 9×10^-8,^ hypergeometric distribution probability test). **i**, Gene ontology analysis reveals that ZIP-5 targets regulate one-carbon metabolism (cysteine and methionine metabolism, purine biosynthesis and glutathione metabolism). **e, f**, Mean ± SEM, *p<0.05, Fisher’s exact test.

Promoter-GFP analysis revealed that ZIP-5 itself is highly expressed in neurons (Fig. 1g), consistent with our prior high-throughput RNA-sequencing of adult neurons and PLM neurons^9^. However, unexpectedly, neither the neuron-expressed a isoform nor the intestine-expressed b isoform of *zip-5* is expressed directly in PLM neurons (Fig. 1g), suggesting a cell non-autonomous role for ZIP-5 activity.

To identify the mechanisms by which ZIP-5 influences axon regeneration, we performed whole-genome expression profiling comparing *daf-2* vs. *daf-2;zip-5* animals (Fig. 1h, Extended Data Fig. 2a, Supplementary tables 2-4). Of the 462 genes upregulated by ZIP-5 at both Day 1 and Day 5, 70% are expressed in neurons (Fig. 1g; p-value = 9×10^-8,^ Extended data fig. 2a, b). ZIP-5 targets regulate the FOXO signaling pathway (Fig. 1i), including *daf-16* (suggesting that ZIP-5 provides positive feedback to its own upstream regulator), *akt-2/AKT2*, the mitochondrial superoxide dismutase *sod-3*, and catalases (*ctl-1, −2, −3*). However, ZIP-5 target genes largely regulate metabolism, including cysteine, methionine, purines, and glutathione (the one-carbon metabolic pathway, Fig. 2a), as well as carbohydrate and lipid metabolism (Fig. 1i, Extended Data Fig. 2b). Unlike DAF-16, whose gene expression differs significantly between neurons and other tissues^9^, the classes of genes that ZIP-5 regulates are consistent between neurons and the whole body, underscoring its role in regulation of metabolism.

**Figure 2.**
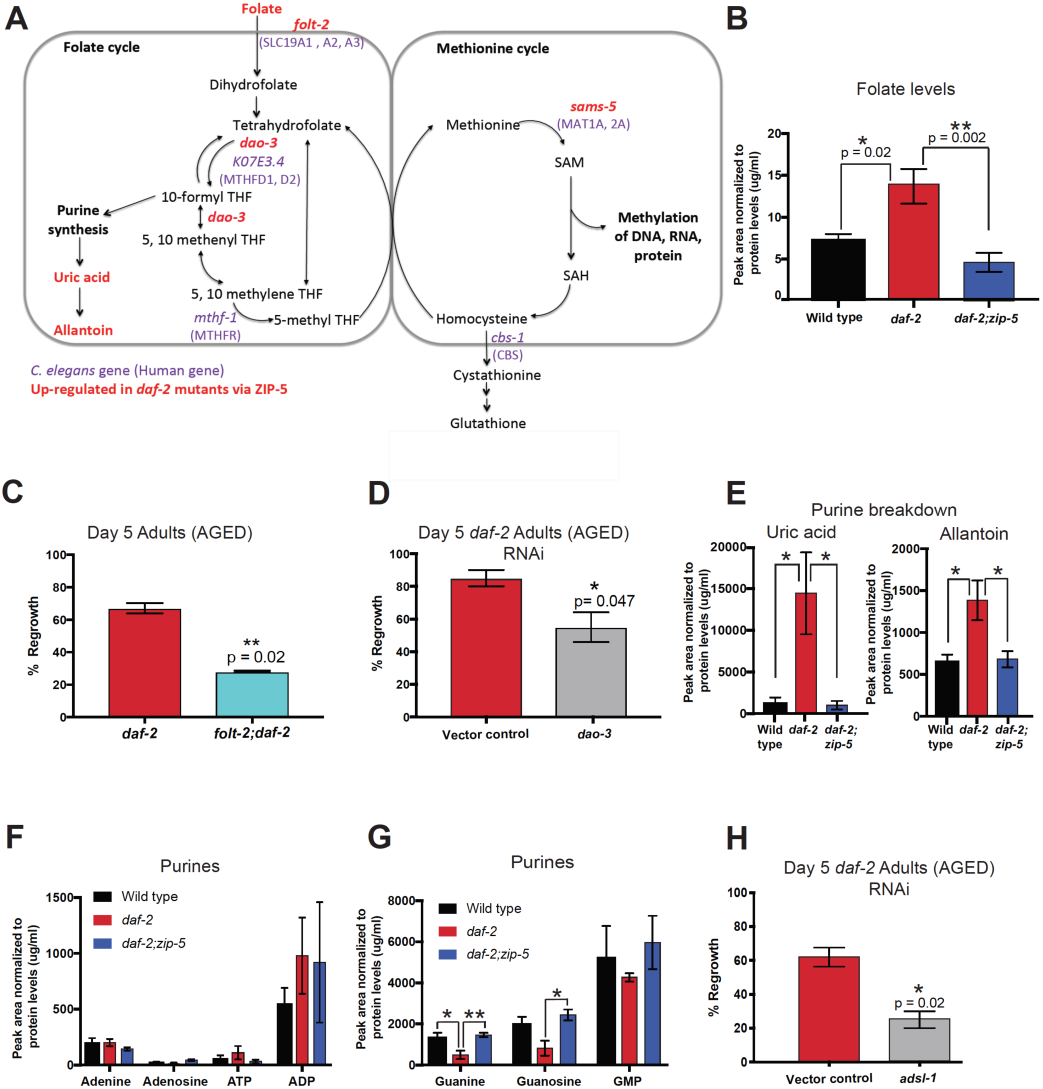
Folate and purine metabolism are upregulated in *daf-2* mutants and is required for the regeneration of aging axons. a, The highly conserved folate cycle feeds metabolites into the methionine and glutathione cycles. b, Folate levels are higher in ***daf-2*** mutants compared to wild type as well as ***daf-2;zip-5*** mutants. c, Knocking down the folate transporter ***folt-2*** reduces age-dependent axon regeneration in *daf-2* mutants. d, RNAi knockdown of *dao-3*, an essential enzyme mediating folate metabolism decreases the regenerative capacity of aging ***daf-2*** mutants. e-g, While absolute levels of most purine metabolites are not altered across wild type, ***daf-2*** and ***daf-2;zip-5*** mutants, the products of purine metabolism (uric acid and allantoin) are higher in *daf-2* mutants. h, Knocking down purine biosynthesis in ***daf-2*** mutants reduces the regeneration of aging axons. b-h, Mean ± SEM, *p<0.05, **p<0.01, b, e, f, g, One-way ANOVA with Tukey’s multiple comparisons test. c, d, h, Fisher’s exact test.

One-carbon metabolism is mediated by the co-factor folate and is highly conserved, functioning in all eukaryotic cells to support multiple physiological processes. Essential molecules produced via the one-carbon pathway include purines, which are used to generate DNA and RNA, SAM/SAH (used in cellular methylation reactions), and glutathione, which maintains cellular redox status^5^ (Fig. 2a). Metabolomic analysis of *daf-2* mutants revealed increased folate levels, while knocking out *zip-5* in *daf-2* mutants reduced folate levels to wild-type levels (Fig. 2b). To test whether increased folate levels are required for *daf-2’s* enhanced axon regeneration, we knocked down the *folt-2* folate transporter, and observed significantly reduced regeneration of aging *daf-2* mutants (Fig. 2c, Extended data fig. 3a). Similarly, levels of the human methylenetetrahydrofolate dehydrogenase (MTHFD1) homolog *dao-3* are higher in *daf-2* mutants and are regulated in a ZIP-5-dependent manner (Supplementary Table 4), and knocking down *dao-3* in *daf-2* mutants also reduced their regenerative capacity at Day 5 of adulthood (Fig. 2d). Products of purine catabolism such as uric acid and allantoin are significantly higher in *daf-2* mutants compared to wild type or *daf-2;zip-5* mutants (Fig. 2e), despite no change in absolute purine levels (Fig. 2f,g), suggesting that *daf-2* mutants utilize purines at a faster rate than wild-type animals and *daf-2;zip-5* mutants.

**Figure 3.**
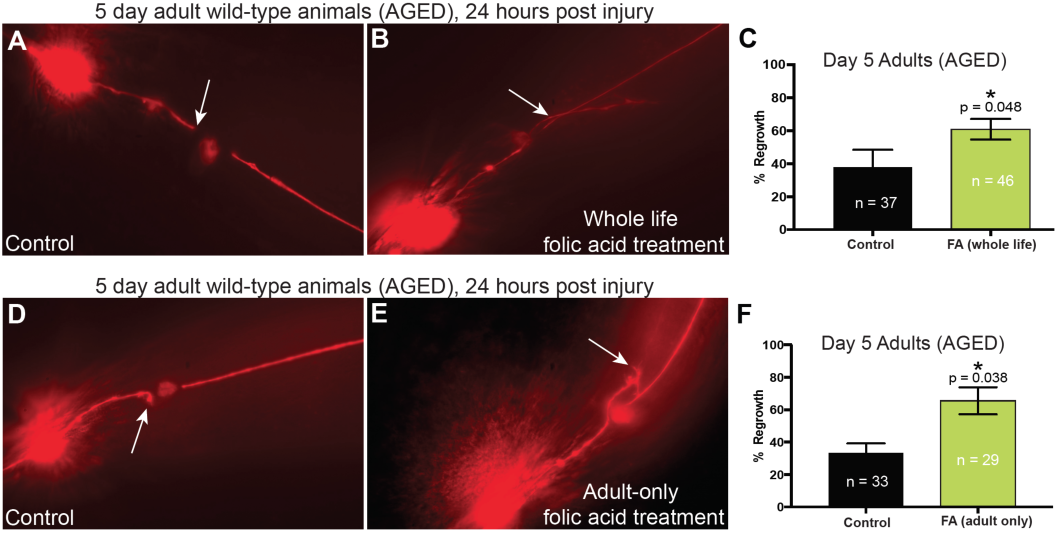
Dietary supplementation with folic acid improves the regeneration of aging *C. elegans* neurons. a-c, Worms treated with folic acid for their whole lives had improved regeneration of mechanosensory neurons at Day 5 of adulthood. d-f, Dietary supplementation with folic acid exclusively in adulthood resulted in improved axonal regeneration in aging animals. c, f, Mean ± SEM, *p<0.05, Fisher’s exact test.

We wondered whether this accelerated purine metabolism in *daf-2* mutants is required for improving age dependent axonal repair. Indeed, knocking down purine biosynthesis in *daf-2* mutants by reducing levels of adenylosuccinate lyase *adsl-1* significantly decreased the regeneration of aging axons (Fig. 2h). Other metabolites of the one-carbon pathway such as methionine, homocysteine, cystathionine, cysteine, glutathione and glutathione disulfide levels are unchanged across wild type, *daf-2*, and *daf-2;zip-5* mutants (Extended data fig. 3b-g). Consistent with these data, knocking down the ZIP-5 target gene *sams-5* (S-Adenosyl Methionine Synthetase) that regulates the levels of these metabolites, does not affect regeneration in aging *daf-2* mutants (Extended data fig. 3h). Thus, the enhanced one-carbon metabolism of *daf-2* mutants likely exerts its regenerative effects by altering the folate cycle and purine metabolism.

To examine whether exogenously boosting one-carbon metabolism could improve axonal regeneration in aging wild-type *C. elegans*, we supplemented their diet with 25 μM folic acid either throughout life (Fig. 3a-c) or only in adulthood (Fig. 3d-f). In both cases, folic acid treatment significantly improved PLM axon regeneration of aging wild-type animals, suggesting that dietary interventions can extend the regenerative capacity of aging axons, independent of larval stage growth and development effects. These data are consistent with the fact that ZIP-5 activity improves PLM regeneration in *daf-2* mutants by upregulating one-carbon metabolism genes in a cell non-autonomous manner, which in turn affects global metabolism.

Human liver, mouse lung and heart, and zebrafish fin tissue exhibit reduced regeneration with age^1, 21, 22^, and older peripheral neurons also dramatically decline in their ability to regrow injured axons^23^. Given the conserved role of folate metabolism in regulating neuronal health in aging animals, we wondered whether boosting folate metabolism could improve the regeneration of aging tissues in vertebrates.

Zebrafish fin tissue has the extraordinary ability to fully regenerate upon amputation. Axonal regeneration into the healing tissue is a prerequisite for fin regrowth in young animals^15^; thus, improving axonal regeneration could promote fin tissue repair in aging animals. Several signaling pathways, such as WNT, FGF, BMP, and insulin signaling play a conserved role in regulating injury-induced regeneration in *C. elegans* axons and zebrafish fins in young animals^3, 26^; therefore, regulators of axonal and tissue repair in aging animals may be similarly conserved. We analyzed previously published high-throughput RNA sequencing data of regenerating zebrafish fins^10^ and found that enzymes regulating folate metabolism are upregulated in zebrafish fins four days post-amputation (Fig. 4a-b, Supplementary Table 6). This mirrors our observation that *daf-2* animals with improved age-dependent axon regeneration specifically have higher activity of folate, but not methionine and glutathione metabolic cycles, underscoring the importance of folate metabolism in regulating cellular repair.

**Figure 4.**
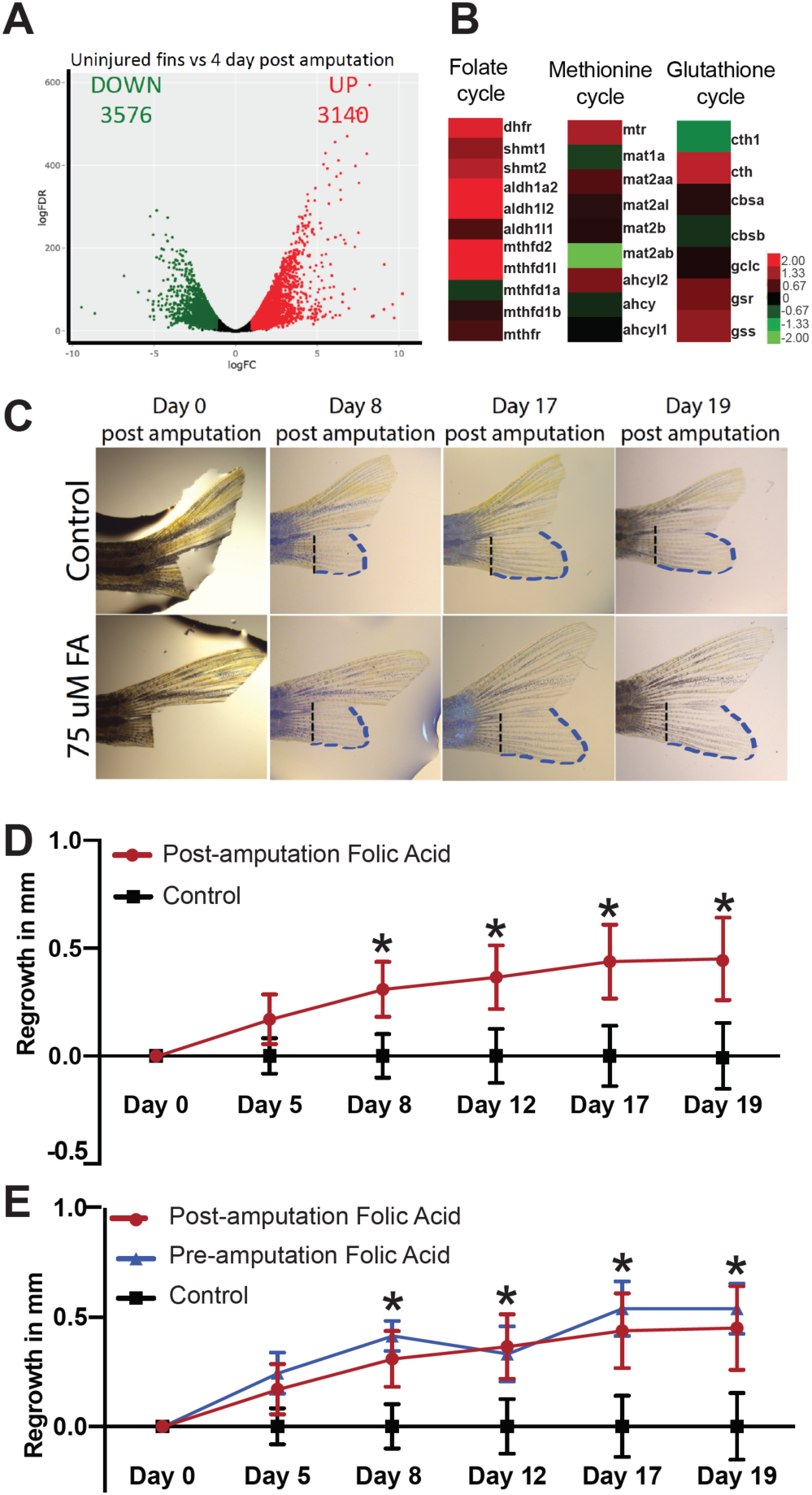
Dietary supplementation with folic acid improves the regeneration of zebrafish fin tissue in aging animals. a,b, High throughput RNA-sequencing of regenerating zebrafish fins identifies 3140 up-regulated transcripts, including enzymes regulating folate metabolism, but not methionine and glutathione metabolism. c,d, Supplementing the diet of aging zebrafish with folic acid after amputation improves fin regeneration. e, Folic acid supplementation after amputation improves fin regeneration to the same extent as folic acid pre-treatment before amputation. d, e, Mean ± SEM, *p<0.05, Two-way ANOVA.

Remarkably, we find that supplementing the diet of aging zebrafish with 75 μM folic acid, even when treated after amputation, improved the rate of fin regeneration (Fig. 4c,d), and pre-treatment with folic acid before amputation did not confer additional benefit towards fin regeneration (Fig 4e). The enhancement of fin regeneration commenced by Day 8 and remained higher until Day 19 post-amputation. Thus, dietary supplementation of folic acid plays a conserved role in improving the regeneration of injured axons and tissues in aging animals. Given that molecular mechanisms required to regenerate old neurons are distinct from those required to repair young neurons, folic acid supplementation represents a unique, non-invasive therapy to repair axons in older animals.

Aging is associated with decreased function of the one-carbon metabolic pathway in humans^20^. Plasma folate levels decrease with age, with a concomitant increase in homocysteine levels^20^. Hyperhomocysteinemia increases the risk for cognitive impairment, dementia, Parkinson’s disease, and stroke. Thus, boosting folate metabolism in elderly humans could have neuroprotective and pro-regenerative effects similar to what we observe here. Consistent with this idea, elderly humans who have high purine catabolism (measured by high uric acid levels) have improved neuronal health and reduced incidence of dementia later in life^6^. Conversely, patients with Parkinson’s, Alzheimer’s disease, or Multiple Sclerosis display lower purine catabolism in the brain and blood^4, 12, 24^. Thus, increased folate and purine metabolism may play a conserved role in slowing neuronal aging. Our data suggest that maintaining folate metabolism with age boosts the regenerative ability of aging neurons and tissues across species. Unbiased bioinformatics analysis predicted that the human homologs of *C. elegans* ZIP-5 targets, including those regulating one-carbon metabolism, play a role in neurodegenerative diseases (Supplementary Table 5). In fact, one-carbon metabolism is upregulated in multiple longevity mutants^14^, and folic acid supplementation in wild-type animals increases lifespan^18^, suggesting a central role for folate metabolism in mediating longevity as well as repair. Our work suggests that dietary folate supplementation should be further developed as a novel approach to promote healthy brain aging, and also as a non-invasive treatment for patients with nerve trauma or neurodegenerative diseases associated with aging.

## Methods

### Strains and worm cultivation

*zip-5(gk646*) and *daf-2 (e1370)* were provided by the CGC which is funded by NIH Office of Research Infrastructure Programs (P40 OD010440). ZB4065 (*Pmec-4::mCherry)* was obtained from Monica Driscoll, Rutgers University. These strains were crossed to generate *CQ461 (daf-2(e1370);Pmec-4::mCherry)* and *CQ501 (daf-2 (e1370);zip-5(gk646);Pmec-4::mCherry). Pzip-5a::GFP* was obtained from Ian Hope, University of Leeds^19^ and was crossed with *Pmec-4::mCherry* to obtain CQ472 (*Pzip-5a::GFP; Pmec-4::mCherry;). Pzip-5b::GFP* was injected into *Pmec-4::mCherry* worms at 10ng/ul along with *Pmyo-2::mCherry* 1ng/ul) to generate CQ519 (*Pzip-5b::GFP;Pmec-4::mcherry;*). Worms carrying a genomic deletion of *folt-2* were generated at Knudra Transgenics (Murray, UT) and were crossed to generate CQ560 (*folt-2(knu-32);Pmec-4::mCherry*).

### Axon regeneration assays

In vivo laser microsurgery of PLM neurons was performed as described^9^. All experiments were repeated 2-3 times. Number of animals used were as follows: Day 1 *daf-2* (n=22) vs. *daf-2;zip-5* (n=23); Day 5 *daf-2* (n=30); vs. *daf-2;zip-5* (n=25); *daf-2* (n=21) vs. *folt-2;daf-2* (n=18); vector control (n=20) vs. *dao-3* RNAi (n=22); vector control (n=21) vs. *adsl-1* RNAi (n=22); vector control (n=25) vs. *sams-5* RNAi (n=27); control (n=37) vs. whole-life folic acid treatment (n=46); control (n=33) vs. adult-only folic acid treatment (n=29).

### Folic acid supplementation

200 ml of 25uM folic acid (Sigma, catalog number: F8798) was added to HG plates seeded with OP50 a few minutes before worm transfer. Worms were bleached onto folic acid-treated plates and transferred to fresh folic acid-treated plates at the L4 stage, or they were transferred onto folic acid-treated plates at Day 1 of adulthood. Adult worms were transferred to fresh folic acid-treated plates daily.

### Microarrays

Day 1 and Day 5 adult *daf-2* and *daf-2;zip-5* worms were collected by crushing in liquid N2. RNA was extracted using the standard Trizol/chloroform/isopropanol method and purified on an RNEasy column (Qiagen). RNA was labeled with Cy3- or Cy5-CTP (Perkin Elmer) using the Agilent Low RNA Input Linear Amplification Kit. cRNA was hybridized on C. elegans (V2) Gene Expression Microarray 4×44K (Agilent) as previously described^13^. Four biological replicates for Day 1 worms and five biological replicates for Day 5 worms were hybridized.

### Microarray analysis

We performed data filtering to eliminate spots whose intensity was not above background, and to identify genes with > 60% good data. The filtered gene list was used for subsequent analysis. To identify genes regulated by ZIP-5 in *daf-2* mutants, we used Significance Analysis of Microarrays (SAM) at 2% FDR to compare gene expression in *daf-2* vs. *daf-2;zip-5* mutants^25^. Data are available at PUMAdb (http://puma.princeton.edu). Genes upregulated in *daf-2* mutants but not in *daf-2;zip-5* mutants were termed ‘ZIP-5 targets’. Whole body ZIP-5 targets were filtered through our list of neuron-expressed genes^9^ to identify neuronal ZIP-5 targets. *The* ***D****atabase for* ***A****nnotation*, ***V****isualization and****I****ntegrated* ***D****iscovery (****DAVID*** *) v6.8 or the* Gene Ontology enRIchment anaLysis and visuaLizAtion tool (GOrilla) was used to identify the enrichment of Gene Ontology terms in gene lists.

### Microscopy

Day 1 and 5 *Pzip-5a::GFP; Pmec-4::mCherry* and *Pzip-5b::GFP; Pmec-4::mCherry* transgenics were imaged on a Nikon TiE microscope using z-stacks. Maximum Intensity Projections made using Nikon *NIS-Elements* are shown.

### Metabolomics

Tissue from wild type, *daf-2* and *daf-2;zip-5* worms (five biological replicates) was obtained by crushing in liquid N2. This was followed by LC-MS analysis as previously described^7^.

### Zebrafish fin regeneration assays

Zebrafish were reared in the Burdine Laboratory Fish Facility. All procedures were performed in accordance with Princeton University’s Institutional Animal Care and Use Committee. Two-year-old zebrafish were anesthetized using tricaine (MS222), and the ventral half of their caudal fin was amputated using a razor blade. Half of the animals were then transferred into 75 mM folic acid (Sigma, catalog number: F8798) dissolved in system water. Fish were transferred into fresh folic acid solution daily. Number of animals used were as follows: Control (n=24), post-amputation folic acid treated (n=23) and pre-amputation folic acid treated (n=8).

### Regenerating fin transcriptome analysis

High throughput RNA-sequencing data comparing regenerating vs. control adult zebrafish caudal fins^10^ was accessed at the zfRegeneration database^16^ (http://www.zfregeneration.org/). This database was used for differential expression analysis and volcano plot generation.

## Supporting information

Supplementary Table 1 - DAF-16 neuronal targets and larval regeneration genes

Supplementary Table 2 - Day1 SAM results

Supplementary Table 3 - Day5 SAM results

Supplementary Table 4 - ZIP-5 targets

Supplementary Table 5 - Human orthologs of ZIP-5 targets

Supplementary Table 6 - Zebrafish fin regeneration transcriptome

## References

1. Anchelin, M., Murcia, L., Alcaraz-Perez, F., Garcia-Navarro, E. M. & Cayuela, M. L. Behaviour of telomere and telomerase during aging and regeneration in zebrafish. PLoS One 6, e16955 (2011).

2. Byrne, A. B. et al. Insulin/IGF1 signaling inhibits age-dependent axon regeneration. Neuron 81, 561–573 (2014).

3. Chen, L. et al. Axon regeneration pathways identified by systematic genetic screening in C. elegans. Neuron 71, 1043–1057 (2011).

4. Church, W. H. & Ward, V. L. Uric acid is reduced in the substantia nigra in Parkinson’s disease: effect on dopamine oxidation. Brain Res. Bull. 33, 419–425 (1994).

5. Ducker, G. S. & Rabinowitz, J. D. One-Carbon Metabolism in Health and Disease. Cell. Metab. 25, 27–42 (2017).

6. Euser, S. M., Hofman, A., Westendorp, R. G. & Breteler, M. M. Serum uric acid and cognitive function and dementia. Brain 132, 377–382 (2009).

7. Fenton, A.R., Janowitz, H.N., McReynolds, M.R., Wang, W. & Hanna-Rose, W. A Caenorhabditis elegans model of adenylosuccinate lyase deficiency reveals neuromuscular and reproductive phenotypes of distinct etiology. https://doi.org/10.1101/181719 (2017).

8. Hwang, I. et al. FOXO protects against age-progressive axonal degeneration. Aging Cell. 17, 10.1111/acel.12701. Epub 2017 Nov 26 (2018).

9. Kaletsky, R. et al. The C. elegans adult neuronal IIS/FOXO transcriptome reveals adult phenotype regulators. Nature 529, 92–96 (2016).

10. Kang, J. et al. Modulation of tissue repair by regeneration enhancer elements. Nature 532, 201–206 (2016).

11. Kauffman, A. L., Ashraf, J. M., Corces-Zimmerman, M. R., Landis, J. N. & Murphy, C. T. Insulin signaling and dietary restriction differentially influence the decline of learning and memory with age. PLoS Biol. 8, e1000372 (2010).

12. Kim, T. S. et al. Decreased plasma antioxidants in patients with Alzheimer’s disease. Int. J. Geriatr. Psychiatry 21, 344–348 (2006).

13. Lakhina, V. et al. Genome-wide Functional Analysis of CREB/Long-Term Memory-Dependent Transcription Reveals Distinct Basal and Memory Gene Expression Programs. Neuron 85, 330–345 (2015).

14. Liu, Y. J. et al. Glycine promotes longevity in Caenorhabditis elegans in a methionine cycle-dependent fashion. PLoS Genet. 15, e1007633 (2019).

15. Meda, F. et al. Nerves Control Redox Levels in Mature Tissues Through Schwann Cells and Hedgehog Signaling. Antioxid. Redox Signal. 24, 299–311 (2016).

16. Nieto-Arellano, R. & Sanchez-Iranzo, H. zfRegeneration: a database for gene expression profiling during regeneration. Bioinformatics 35, 703–705 (2019).

17. Ramos, M. I. et al. Low folate status is associated with impaired cognitive function and dementia in the Sacramento Area Latino Study on Aging. Am. J. Clin. Nutr. 82, 1346–1352 (2005).

18. Rathor, L., Akhoon, B. A., Pandey, S., Srivastava, S. & Pandey, R. Folic acid supplementation at lower doses increases oxidative stress resistance and longevity in Caenorhabditis elegans. Age (Dordr) 37, 113-015–9850-5. Epub 2015 Nov 6 (2015).

19. Reece-Hoyes, J. S. et al. Insight into transcription factor gene duplication from Caenorhabditis elegans Promoterome-driven expression patterns. BMC Genomics 8, 27-2164–8-27 (2007).

20. Selhub, J., Jacques, P. F., Wilson, P. W., Rush, D. & Rosenberg, I. H. Vitamin status and intake as primary determinants of homocysteinemia in an elderly population. JAMA 270, 2693–2698 (1993).

21. Shirabe, K. et al. Human early liver regeneration after hepatectomy in patients with hepatocellular carcinoma: special reference to age. Scand. J. Surg. 102, 101–105 (2013).

22. Sousounis, K., Baddour, J. A. & Tsonis, P. A. Aging and regeneration in vertebrates. Curr. Top. Dev. Biol. 108, 217–246 (2014).

23. Tanaka, K. & Webster, H. D. Myelinated fiber regeneration after crush injury is retarded in sciatic nerves of aging mice. J. Comp. Neurol. 308, 180–187 (1991).

24. Toncev, G., Milicic, B., Toncev, S. & Samardzic, G. Serum uric acid levels in multiple sclerosis patients correlate with activity of disease and blood-brain barrier dysfunction. Eur. J. Neurol. 9, 221–226 (2002).

25. Tusher, V. G., Tibshirani, R. & Chu, G. Significance analysis of microarrays applied to the ionizing radiation response. Proc. Natl. Acad. Sci. U. S. A. 98, 5116–5121 (2001).

26. Wehner, D. & Weidinger, G. Signaling networks organizing regenerative growth of the zebrafish fin. Trends Genet. 31, 336–343 (2015).

