## Supplementary Information for "ZIP-5/bZIP transcription factor regulation of folate metabolism is critical for aging axon regeneration"

### Extended Data Fig. 1

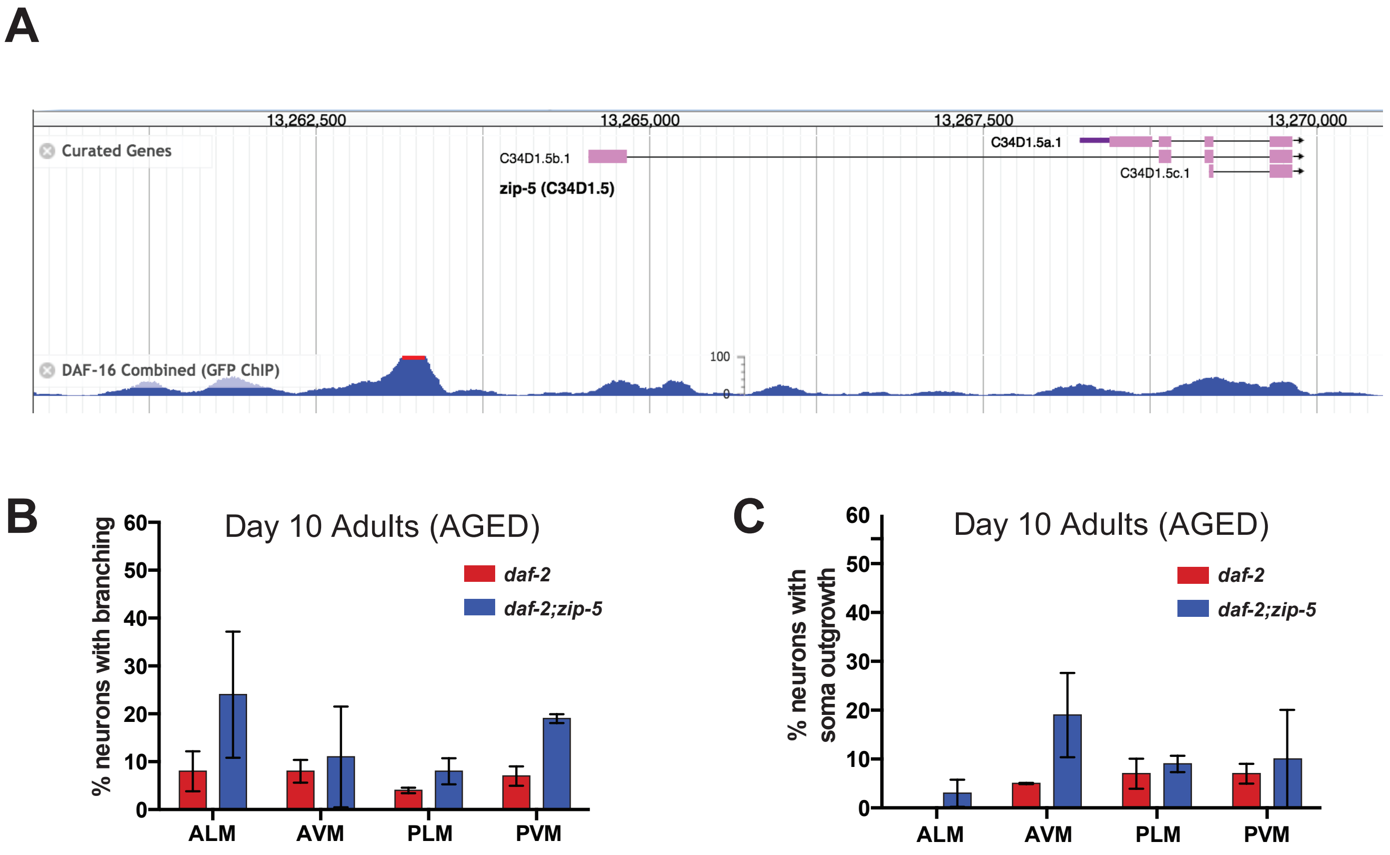


Figure 1: ZIP-5 is a neuron-expressed DAF-16/FOXO target that is not required to prevent age-related morphological defects.

a, Data from the modENCODE project shows that several DAF-16/FOXO binding sites are located upstream to and within the *zip-5* coding region. **b,c,** Reducing *zip-5* levels in *daf-2* mutants does not alter the incidence of age-related morphological defects (axonal branching or soma outgrowth). Mean ± SEM, *$p<0.05$, Fisher's exact test.

### Extended Data Fig. 2

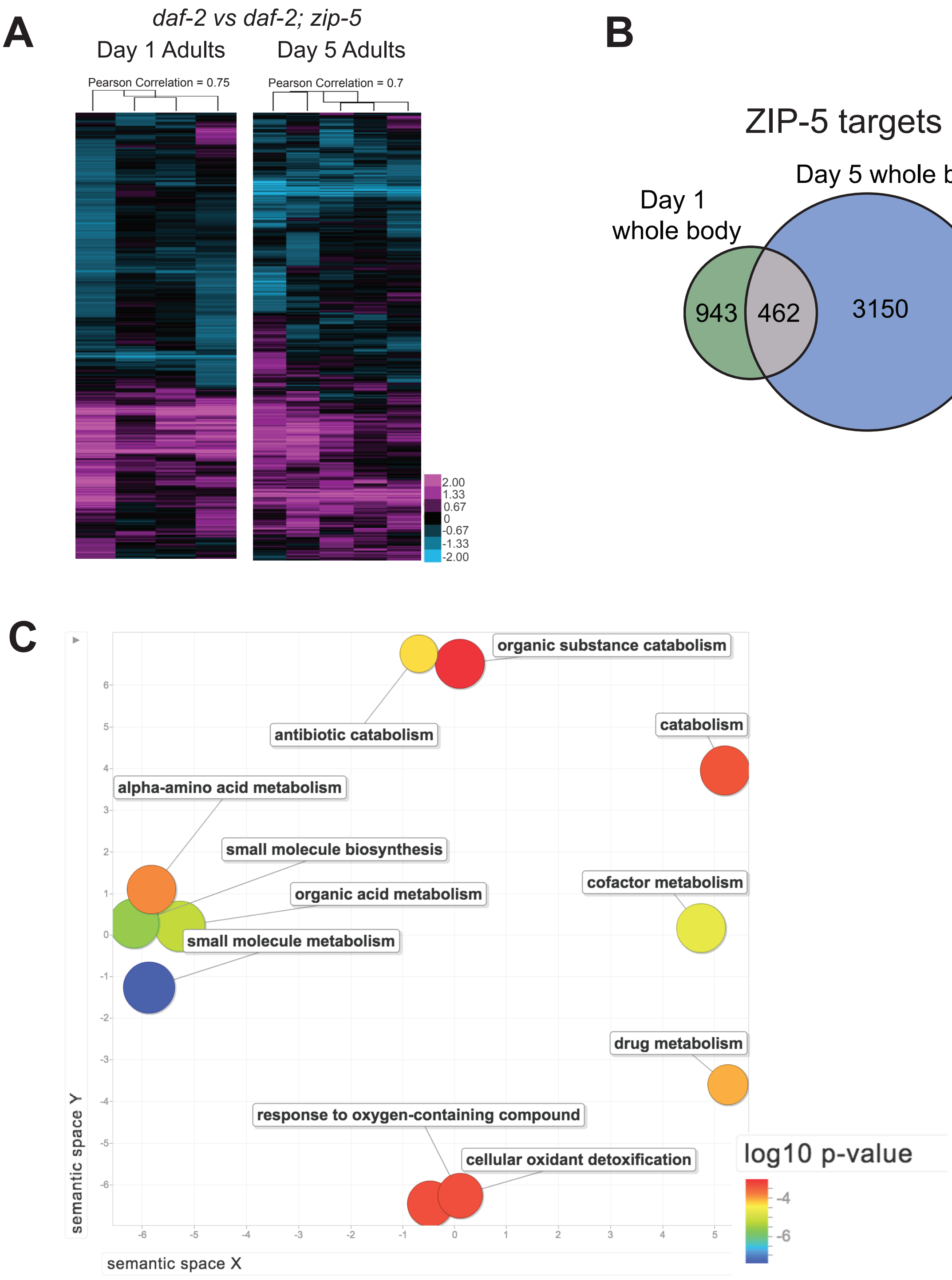


Figure 2: ZIP-5 target genes largely regulate metabolism.

**a, b,** 462 genes are up-regulated by ZIP-5 in both Day 1 and Day 5 adults. **c,** Gene ontology analysis of these 462 genes via REVIGO confirms that these genes are involved in regulating metabolism.

### Extended Data Fig. 3

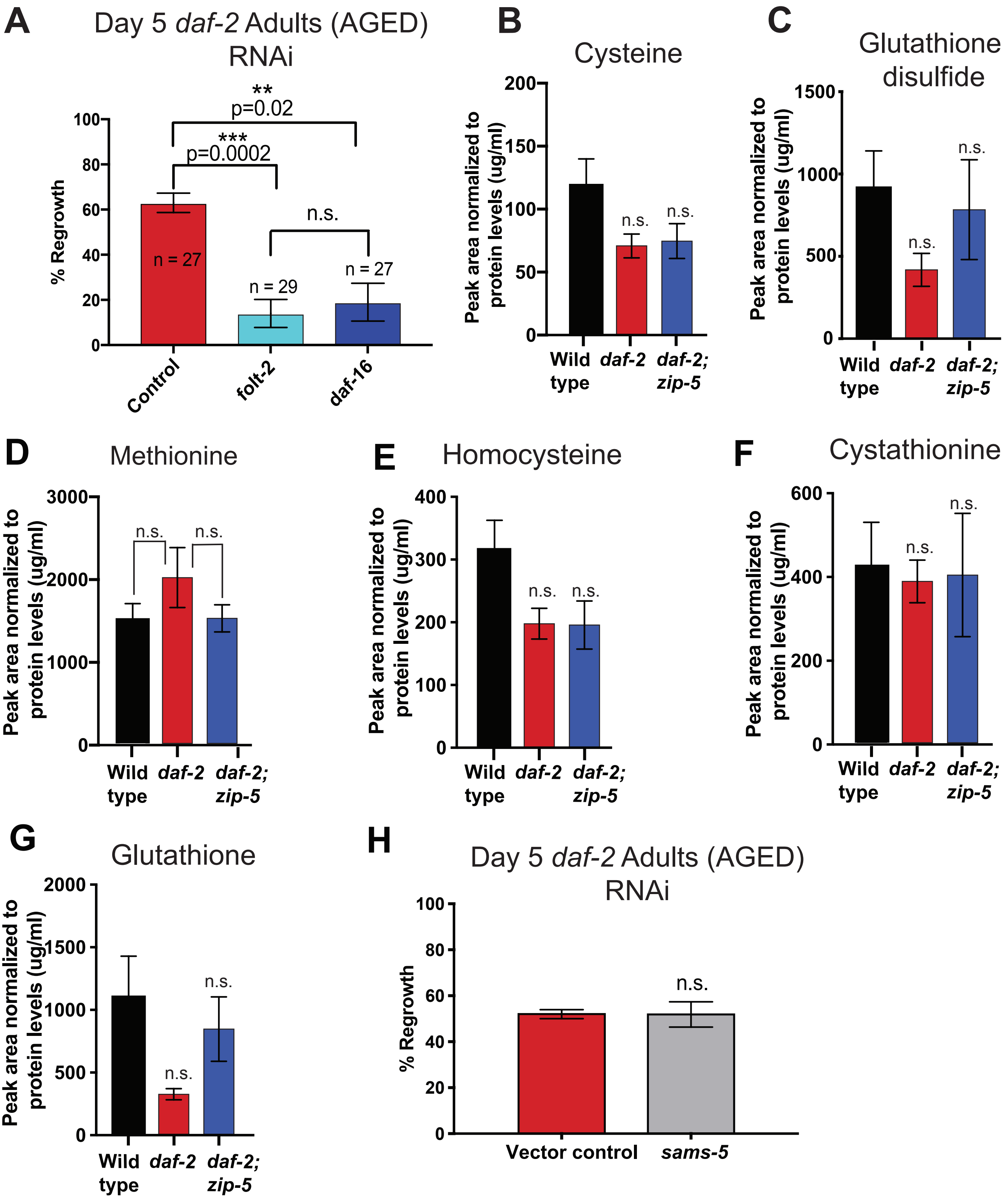

Figure 3: *daf-2* mutants require high folate levels for improved axonal regeneration of aging axons, while other metabolites are unaltered.

**a,** Knocking down the folate transporter *folt-2* via RNAi reduces the regeneration of aging *daf-2* axons, and phenocopies the mutant data shown in Figure 2c. **b-g,** Metabolites of the methionine and glutathione cycles are not altered in wild type, *daf-2* and *daf-2;zip-5* mutants, and altering these metabolites does not influence axon regeneration. **h,** Knocking down the S-Adenosyl Methionine Synthetase *sams-5* does not alter the regenerative capacity of aging *daf-2* axons.

**SUPPLEMENT**

Supplementary tables

Table 1: Genes significantly up- and down- regulated by ZIP-5 in Day 1 adults were identified by significance analysis of microarrays (SAM) analysis of gene expression in *daf-2* vs. *daf-2;zip-5* mutants (FDR = 2%).

Table 2: Genes significantly up- and down- regulated by ZIP-5 in Day 5 aging adults were identified by SAM analysis of gene expression in *daf-2* vs. *daf-2;zip-5* mutants (FDR = 2%).

Table 3: Genes upregulated by ZIP-5 in neurons and the whole body are listed. Gene descriptions were obtained from Wormbase. Gene ontology analysis was performed using DAVID and GOrilla as shown.

Table 4: The human orthologs of ZIP-5 whole body targets were identified using Ortholist 2 (Kim et al., 2018, http://ortholist.shaye-lab.org/). Human orthologs of ZIP-5 targets are enriched for a predicted role in neurodegenerative disease using TargetMine (https://targetmine.mizuguchilab.org/targetmine/begin.do).

Table 5: Transcriptome of a regenerating fin 4 days post amputation (Kang et al., 2016). GO analysis for upregulated genes is shown. Expression changes in one-carbon metabolism genes is listed; genes involved in folate metabolism, but not methionine and glutathione metabolism are upregulated in regenerating zebrafish fins.

**Supplementary References:**

1. Kang, J. *et al*. Modulation of tissue repair by regeneration enhancer elements. *Nature* **532**, 201-206 (2016).

2. Kim, W., Underwood, R. S., Greenwald, I. & Shaye, D. D. OrthoList 2: A New Comparative Genomic Analysis of Human and Caenorhabditis elegans Genes. *Genetics* **210**, 445-461 (2018).
